# Endothelial exosome plays functional role during rickettsial infection

**DOI:** 10.1101/2020.11.16.385740

**Authors:** Yakun Liu, Changcheng Zhou, Zhengchen Su, Qing Chang, Yuan Qiu, Jiani Bei, Angelo Gaitas, Jie Xiao, Alexandra Drelich, Kamil Khanipov, Yang Jin, Georgiy Golovko, Tais B. Saito, Bin Gong

## Abstract

Spotted fever group rickettsioses (SFRs) are devastating human infections. Vascular endothelial cells (ECs) are the primary targets of infection. Edema resulting from EC barrier dysfunction occurs in the brain and lungs in most cases of lethal SFR, but the underlying mechanisms remain unclear. The aim of the study is to explore the potential role of *Rickettsia* (*R*)-infected, EC-derived exosomes (Exos) during infection. Using size-exclusion chromatography (SEC), we purified Exos from conditioned, filtered, bacteria-free media collected from *R*-infected human umbilical vein ECs (HUVECs) (*R*-ECExos) and plasma of *R*-infected mice (*R*-plsExos). We observed that rickettsial infection increases the release of heterogeneous plsExos, but endothelial exosomal size, morphology, and production were not significantly altered following infection. Compared to normal plsExos and ECExos, both *R*-plsExos and *R*-ECExos induced dysfunction of recipient normal brain microvascular Ecs (BMECs). The effect of *R*-plsExos on mouse recipient BMEC barrier function is dose-dependent. The effect of *R*-ECExos on human recipient BMEC barrier function is dependent on exosomal RNA cargo. Next-generation sequencing analysis and stem-loop quantitative reverse transcription PCR (RT-qPCR) validation revealed that *R* infection triggered the selective enrichment of endothelial exosomal mir-23a and mir-30b, which target the endothelial barrier. To our knowledge, this is the first report on the functional role of extracellular vesicles following infection by obligately intracellular bacteria.

**Importance:** Spotted fever group rickettsioses are devastating human infections. Vascular endothelial cells are the primary targets of infection. Edema resulting from endothelial barrier dysfunction occurs in the brain and lungs in most cases of lethal rickettsioses, but the underlying mechanisms remain unclear. The aim of the study is to explore the potential role of *Rickettsia*-infected, endothelial cell-derived exosomes during infection. We observed that rickettsial infection increases the release of heterogeneous plasma Exos, but endothelial exosomal size, morphology, and production were not significantly altered following infection. *Rickettsia*-infected, endothelial cell-derived exosomes induced dysfunction of recipient normal brain microvascular endothelial cells. The effect is dependent on exosomal RNA cargo. Next-generation sequencing analysis revealed that rickettsial infection triggered the selective enrichment of endothelial exosomal mir-23a and mir-30b, which target the endothelial barrier. To our knowledge, this is the first report on the functional role of extracellular vesicles following infection by obligately intracellular bacteria.

## Introduction

Spotted fever group rickettsioses (SFRs) are devastating human infections (1). A licensed vaccine is not available. It forecasted that increased ambient temperatures under conditions of global climate change will lead to more widespread distribution of rickettsioses (2). These arthropod-borne diseases are caused by obligately intracellular bacteria of the genus *Rickettsia* (*R*), including *R. rickettsii* (3, 4) and *R. parkeri* (5-7) that cause Rocky Mountain spotted fever and *R. parkeri* rickettsiosis (8), respectively, in the United States and Latin America; *R. conorii*, the causative agent of Mediterranean spotted fever endemic to southern Europe, North Africa, and India (9); and *R. australis*, which causes Queensland tick typhus in Australia (10). Vascular endothelial cells (ECs) are the primary targets of infection, and EC tropism plays a central role during pathogenesis (1, 3, 11). Edema resulting from EC barrier dysfunction occurs in the brain and lungs in most cases of lethal SFR. Typically, *R* infection is controlled by appropriate broad-spectrum antibiotic therapy if diagnosed early (3, 4). However, *R* infections can cause nonspecific signs and symptoms, rendering early clinical diagnosis difficult (12, 13). Untreated or misdiagnosed *R* infections are frequently associated with severe morbidity and mortality (1, 14-17). A fatality rate as high as 32% has been reported in hospitalized patients with Mediterranean spotted fever (17). Although doxycycline is the antibiotic of choice for *R* infections, it only stops bacteria from reproducing, but does not kill the rickettsiae. Comprehensive understanding of rickettsial pathogenesis is urgently needed for the development of novel therapeutics (7, 16, 18-22).

Eukaryotic cell-to-cell communication is critical for maintaining homeostasis and responding quickly to environmental stimuli (23-51). Besides direct intercellular contact, this communication is often mediated by soluble factors that can convey signals to a large repertoire of responding cells, either locally or remotely. Extracellular vesicles (EVs) transfer functional mediators to neighboring and distant recipient cells (33). EVs are broadly classified into two categories, exosomes (Exos)(50-150 nm) and microvesicles (100-1000 nm), owing to their endocytic or plasma membrane origin (38, 52-66). Exos and microvesicles are also termed as small and large EVs, respectively (55). Exo biogenesis begins with the formation of intraluminal vesicles, the intracellular precursors of Exos, after the inward budding of the membranes of late endosomes (37, 54). Intraluminal vesicles are internalized into a multivesicular body, which transits towards and fuses with the plasma membrane, before releasing intraluminal vesicle into the extracellular environment as Exos (53). An Exo contains many types of biomolecules, including proteins, nucleic acids, and lipids (67). Once bound to the plasma membrane of the recipient cell, Exos can induce functional responses by multiple mechanisms, e.g., activating receptors on recipient cells or releasing their bioactive cargos after internalization (67, 68). In infectious biology, EVs from infected donor cells contain cargos that are associated with the virulence of the pathogen or the activation of host self-defense mechanisms (33-38, 69-71). Evs released from macrophages infected by intracellular bacteria, such as *Mycobacterium tuberculosis* and *Salmonella typhimurium*, have been shown to stimulate a pro-inflammatory response in non-infected macrophages in toll-like receptor-dependent manner (70). Unfortunately, the role(s) of EVs in the pathogenesis of obligately intracellular bacterial infections remains unknown.

Although small noncoding RNA (sncRNA) species (<150□nucleotides) are relatively stable when compared with other RNA molecules, they remain vulnerable to ribonuclease (RNase)-mediated digestion (72). The discovery of extracellular sncRNAs in the blood, despite the abundant presence of RNases, led to the proposal of a scenario in which sncRNAs are encapsulated in EVs (55, 72-74) or form circulating ribonucleoproteins (75, 76). Extracellular RNAs are enriched in sncRNAs (77). A growing number of reports have established that many, if not all, the effects of EVs are mediated by microRNA (52, 55-60, 63) or tRNA fragment (61, 78) cargos, which remain functional to regulate cellular behaviors of the recipient cells (79). Recent studies provide emerging evidence that microRNAs are selectively sorted into EVs independently of their cellular levels (52, 55-62). We reported that *R* infection induces significant upregulation of specific tRNA-derived RNA fragments in host cell, but no global changes of microRNAs in perfusion-rinsed mouse lung tissues was observed (80). Information regarding the potential role of extracellular RNAs during *R* infection is still lacking.

The aim of this study is to explore the potential role of *R*-infected, EC-derived Exos following infection. Using size-exclusion chromatography (SEC), we purified Exos from conditioned, filtered, bacteria-free media collected from *R*-infected human umbilical vein ECs (HUVECs) (*R*-ECExos) and plasma of *R*-infected mice (*R*-plsExos). We observed that, compared to noninfectious normal mouse plsExos and normal HUVEC-derived Exos, both *R*-plsExos and *R*-ECExos induced dysfunction of normal brain microvascular ECs (BMECs). The effect of *R*-plsExos on mouse recipient BMEC barrier function is dose-dependent. The effect of *R*-ECExos on human recipient BMEC barrier function is dependent upon exosomal RNA cargos. Saponin-assisted active exosomal permeabilization pretreatment (81-83) of *R*-ECExos with RNase mitigated the effect of *R*-ECExos on recipient BMEC barrier function. Next-generation sequencing analysis and stem-loop quantitative reverse transcription PCR (RT-qPCR) validation revealed that *R* infection triggered the selective enrichment of endothelial exosomal mir-23a and mir-30b, which target the endothelial barrier.

## Results

### 1. Quality assessment of bacteria-free Exos from the plasma of *R. australis*-infected mice (*R*-plsExos) and the media of *R. parkeri*-infected HUVECs (*R*-ECExos)

Using SEC, we isolated small EVs (50-150 nm) from *R*-infected mouse plasma and HUVECs from culture media; both were passed through two 0.2 µm filters. Quantitative real-time PCR validated that no rickettsial DNA copies were detected in either *R*-plsExos isolated *R. australis*-infected mice infected with 2 LD_50_ of bacteria (80, 84-87) on day 4 post-infection (p.i.), or the *R*-ECExos that were purified 72 hrs p.i. from *R. parkeri*-infected HUVECs (6) using a multiplicity of infection (MOI) of 10 (**Fig. Suppl 1**).

Sizes and morphologies of isolated EVs from mouse plasma and EC culture media, respectively, were initially evaluated using transmission electron microscopy (TEM) (52, 88), or atomic force microscopy (AFM) (89). The images captured using TEM and AFM show particles with typical exosomal morphology (arrowheads in **Fig. 1a**), as published previously (52, 88, 90). Using nanoparticle tracking analysis (NTA), the size distribution of isolated EVs was also confirmed to be in the range of 50 to 150 nm, which is the expected size of Exos (**Figs. 1b and c)**. We also verified the purity of isolated Exos using western immunoblotting to detect traditional exosomal markers as shown in **Fig. 1d** (64, 73, 90).

**Figure 1:**
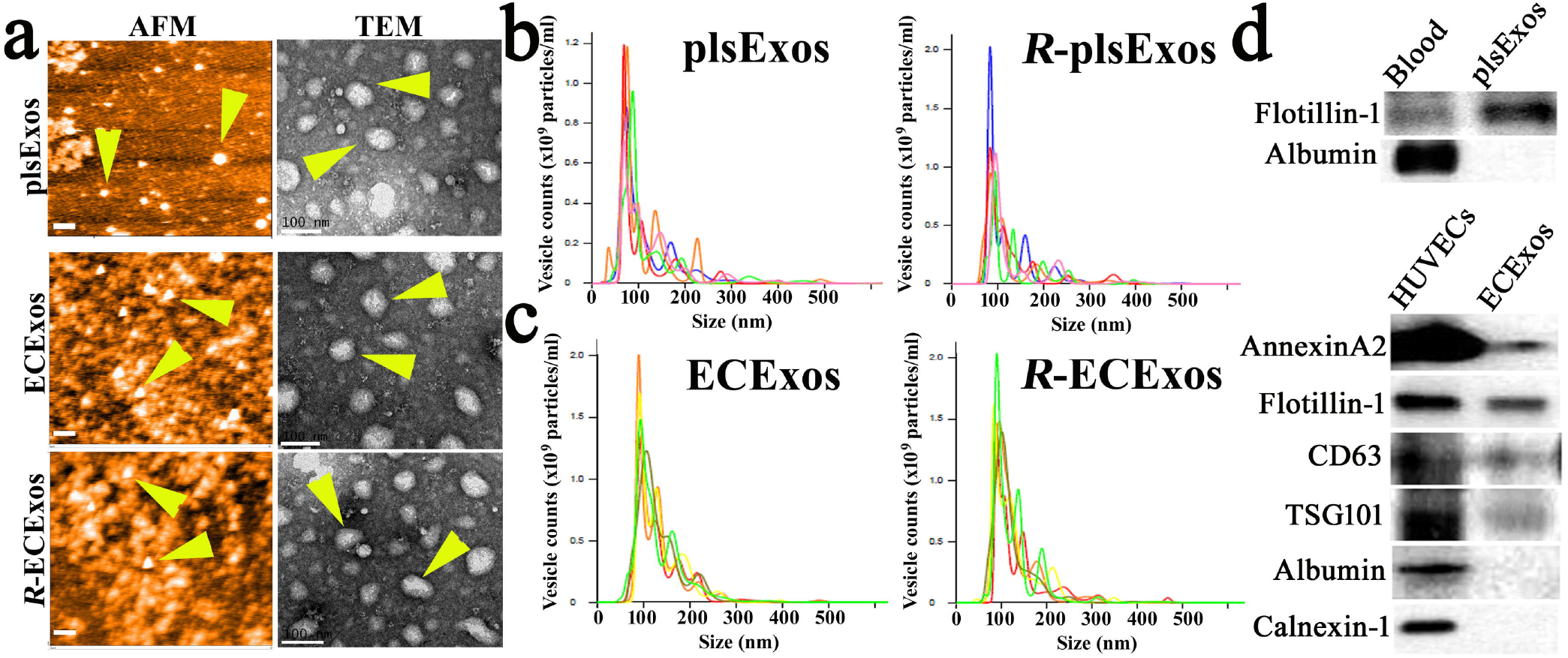
Characterization of plsExos and ECExos after SEC isolation. (**a**) plsExos and ECExos morphologies were verified using atomic force microscopy (AFM)(left panels; scale bars, 200 nm) and transmission electronic microscopy (TEM)(right panels; scale bars, 100 nm). (**b**) and (**c**), the vesicle size distribution of isolated EVs was analyzed using nanoparticle tracking analysis (NTA) (n=5 per group). **d**, Expressions of indicated protein markers in 100 μg proteins of plsExos (upper panel) and ECExos (lower panel) were examined using western immunoblotting.

These data demonstrate that purified EVs from *R*-infected mouse plasma or culture media used in these studies was free of bacteria or bacterial DNA, were intact and did not aggregate, and fell within the expected size the range of Exos.

### 2. Exos are differentially induced and detected in mouse plasma and EC culture media in response to *R* infection

Serum Exos have been identified as being heterogeneous and derived from multiple cell types, including ECs. Using western immunoblotting, we detected EC markers [CD31 and VE-cadherin (91, 92)] in mouse plsExos, as well as markers of other cells (CD45) (**Fig. 2a**), suggesting that the mouse plsExos used in these studies was derived from different types of cells, including ECs.

**Figure 2:**
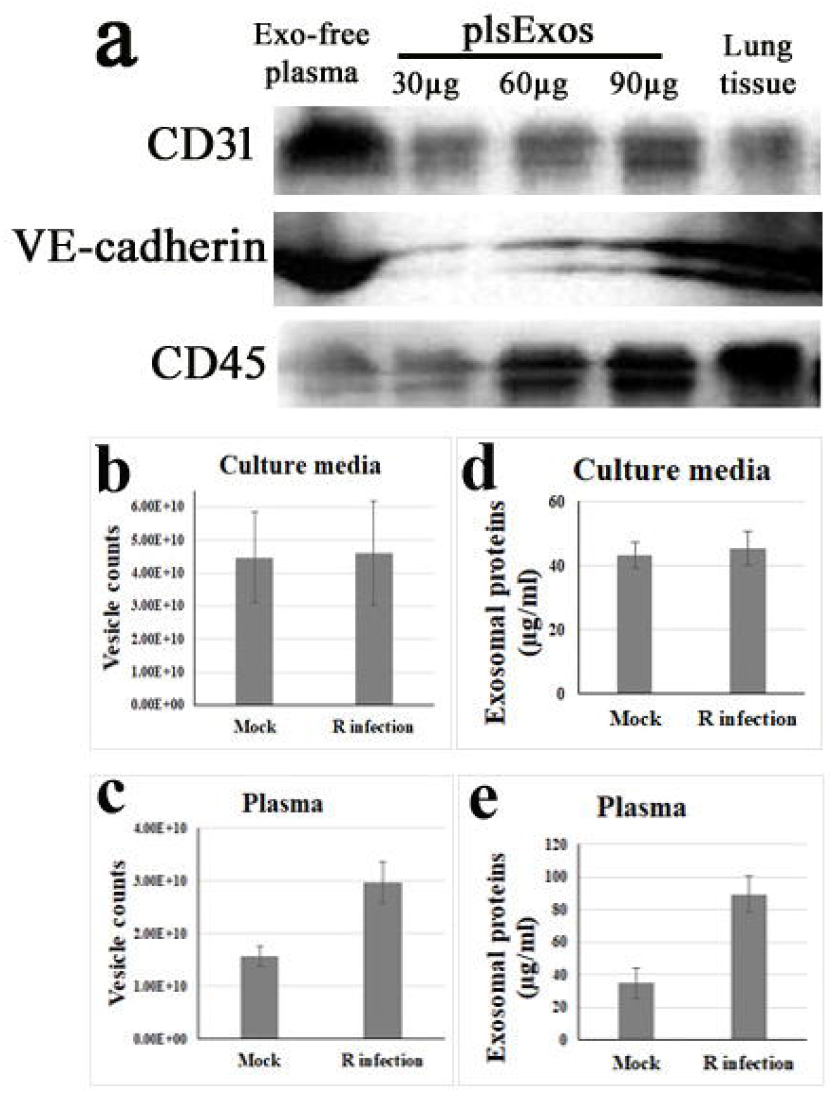
Exos are differentially induced and detected in mouse plasma and EC culture media in response to *R* infection. (**a**) Expression of indicated protein markers (i.e., 30, 60, and 90 μg of plsExos proteins) was examined using western immunoblotting. (**b**) and (**c**), the concentration of plasExos and ECExos was analyzed using NTA (n=5 per group). (**d**) and (**e**), the concentration of exosomal total protein was determined using the microBCA protein assay (n=5 per group).

Exosomal particle counts were measured using NTA, and showed that similar numbers of endothelial Exos are produced by mock and *R* infection groups *in vitro* (**Fig. 2b**). However, the number of mouse *R*-plsExos was upregulated on day 4 p.i. (p=0.02) *in vivo* (**Fig. 2c**). Exos were also assessed using exosomal total protein content (88). As shown in **Figs. 2d and 2e**, the generation of *R*-plsExos was significantly upregulated on day 4 p.i. (p=0.005). Furthermore, we also compared the morphology of EC-derived Exos using TEM and AFM, and demonstrated no significant differences between normal ECExos and *R*-ECExos (**Fig. 1a**).

Collectively, these data suggest that *R* infection increases heterogeneous plsExo release. However, endothelial Exo size, morphology, and production were not significantly altered after infection *in vitro*.

### 3. Recipient cells efficiently take up Exos

ECs are directly exposed to circulating substances and Exos, which are abundant in blood and are taken up by ECs (93, 94). To confirm that ECs take up Exos *in vivo*, we intravenously delivered fluorescent PKH26-labeled plsExos (1 x 10^11^ particles per mouse in 100 µl PBS) to normal mice as described (90). As in **Fig 3a**, co-localization between PKH26 (red) and CD31 (green, a marker of EC lineage) (arrowheads) were identified in multiple organs in mice, which were extensively perfused with PBS at 6 hrs post-injection, prior to fixation. These data suggest that plsExos directly interact with ECs *in vivo*.

**Figure 3:**
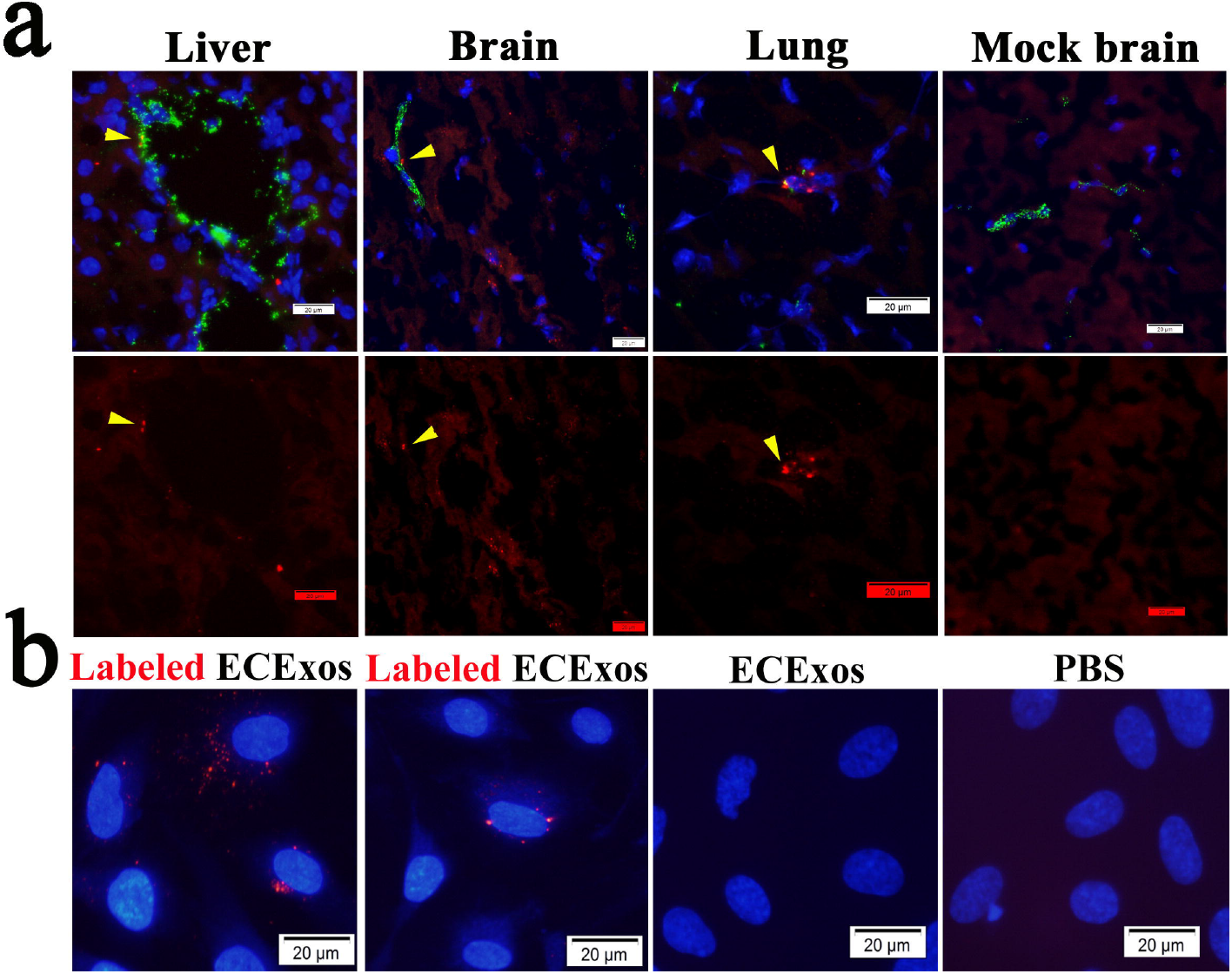
Recipient cells take up Exos. (**a**) Purified plsExos (5 x 10^10^ particles in 50 μL PBS) labeled with PKH26 were administrated to wild-type mice intravenously (n=3). After 4 hrs, organs were dissected for frozen sectioning after euthanasia and perfusion via the right ventricle. Representative immunofluorescent staining of ECs from liver, brain, and lung using an antibody against CD31(an EC marker) is shown. The nuclei were stained with DAPI. Cells with red fluorescence indicate the uptake of PKH26 labeled Exos. Scale bars, 20 µm. (**b**) Purified ECExos were labeled with PKH26 (red) and added to the culture medium of human BMECs (2000 particles per cell) as indicated. Pictures were taken using fluorescence microscopy after 2 hrs of ECExo incubation. Scale bars, 20 µm.

Next, we examined HUVEC-derived ECExo uptake using normal recipient cells (i.e., human BMECs) *in vitro*. PKH26-labeled ECExos and non-labeled controls were added to the cultured human BMECs. As early as 2 hrs after incubation, the uptake of PKH26-labeled ECExos by BMECs was visualized using fluorescence microscopy (**Fig. 3b**).

These data suggest that vascular ECs efficiently take up Exos in our models.

### 4. Effect of mouse *R*-plsExos on normal mouse recipient ECs

We next sought to evaluate the potential effect of *R*-plsExo on normal recipient ECs during *R* infection. Using SEC, *R*-plsExos from a mouse that was intravenously infected with a 2 LD_50_ dose of *R. australis* (86, 87) were isolated. Normal mouse recipient BMECs were treated with normal plsExos or *R*-plsExos at different doses (i.e., 8, 40, or 160 pg Exos/per cell) for 72 hrs before measuring the transendothelial electrical resistance (TEER), an indicator for endothelial paracellular barrier function (95). We found that, compared to normal mouse plsExos, mouse *R*-plsExos derived on day 4 p.i. reduced the TEER in normal recipient ECs in a dose-dependent manner (**Fig. 4a**).

**Figure 4:**
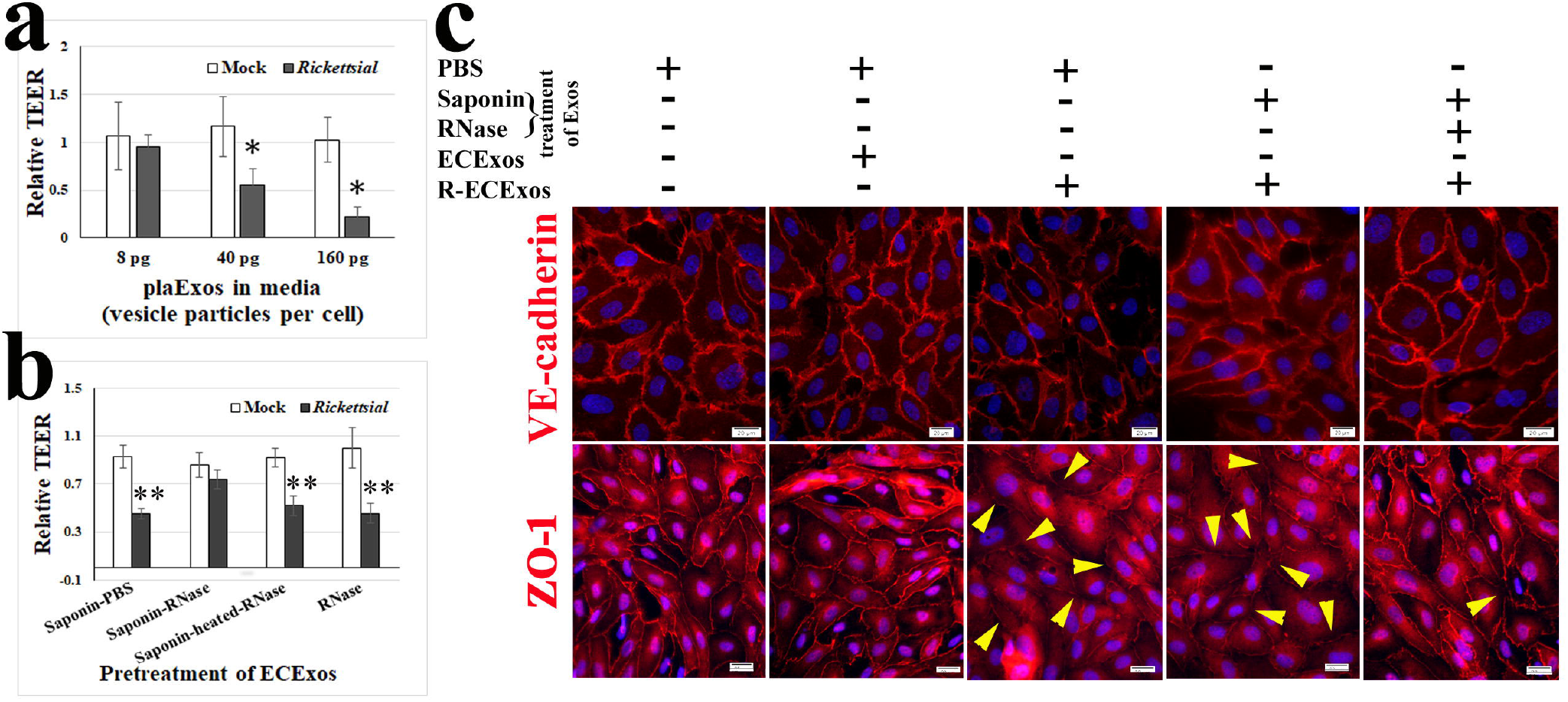
Effect of *R*-plsExos or *R*-ECExos on normal recipient ECs. (**a**) The transendothelial electrical resistance (TEER) values of normal mouse recipient BMECs was measured after treatment with normal plsExos (mock) or R-plsExos at 8, 40, or 160 pg Exos per cell for 72 hrs. *, P < 0.05. (**b**) The TEER values of normal mouse recipient BMECs was measured after a 72 hr-treatment with normal plsExos (mock) or *R*-plsExos, which were pretreated with 20 µg/mL ribonuclease (RNase) in the presence or absence of 0.1% saponin. **, P < 0.01. (**c**) Immunofluorescence staining of tight junctional protein ZO-1 and adherens junctional protein VE-cadherin in normal human recipient BMECs that were treated with different Exos for 72 hrs. Scale bars, 20 μm.

This evidence suggests that mouse *R*-plsExos induce dysfunction in normal recipient ECs in a dose-dependent manner.

### 5. Human *R*-ECExos induced dysfunction of normal human recipient ECs in an exosomal RNA-dependent manner

Endothelial markers were detected in plsExos (**Fig. 2a**). Given that ECs are the major target cells during *R* infection, HUVEC-derived ECExos were used to explore their effect on normal human recipient BMEC function.

It was first found that, compared with normal (mock) ECExos (40 pg vesicle particles), *R*-ECExos (40 pg vesicle particles) reduced TEER in normal recipient BMECs (**Fig. 4b**). Furthermore, *R*-ECExos (40 pg particles/cell) weakened the tight junctional protein ZO-1 (arrowheads in **Fig. 4c**) of normal recipient BMECs. However, we did not observe remarkable alteration of adherens junctional protein VE-cadherin (**Fig. 4c**).

Exos contain many types of biomolecules, including proteins and nucleic acids, which contribute to disease pathogenesis (68). Active encapsulation techniques have been widely employed in the field of EV research, showing no significant impairment of exosomal constitution, integrity, and functionality (81-83). To identify the functional exosomal cargos during *R* infection, we employed saponin-assisted active permeabilization (81-83) to pretreat exosomal cargos with 20 µg/mL RNase in the presence of 0.1 mg/ml saponin. Such pretreatment of *R*-ECExos mitigated the effect on TEER in normal recipient BMECs, compared to RNase in the absence of permeabilization or heat-treated RNase in the presence of saponin (**Fig. 4b**). Similar pretreatment of *R*-ECExos with RNase in the presence of saponin also impaired the tight junctional protein ZO-1 in recipient BMECs (**Fig. 4c**).

These data suggest that *R*-ECExos can induce normal recipient EC barrier dysfunction in an exosomal RNA cargo-dependent manner.

### *6. R* infection upregulates exosomal mir-23a and mir-30b

EV RNA cargo mostly consists of sncRNAs, mainly microRNAs and tRNA-derived fragments (61, 77, 78). A growing number of reports have established that many effects of EVs are mediated by microRNAs (52, 55-60, 63). We characterized the exosomal microRNA cargo using next-generation sequencing (**Fig. 5a**). RNAs were isolated from Exos released from HUVECs infected with *R. parkeri* (at 10 MOI) for 72 hrs or mock-infected. *R. parkeri* is a BSL-2 pathogen, facilitating us to do mechanism studies. There were no differences in total sncRNAs (<150 nucleotides) obtained per Exo between normal ECExo and *R*-ECExo. Seventy-two hours after *R* infection, mir-23a and mir-30b exhibited the greatest induction of expression in *R*-ECExos, reaching 7.69-fold and 3.04-fold increases compared to controls, respectively (**Fig. 5b**).

**Figure 5:**
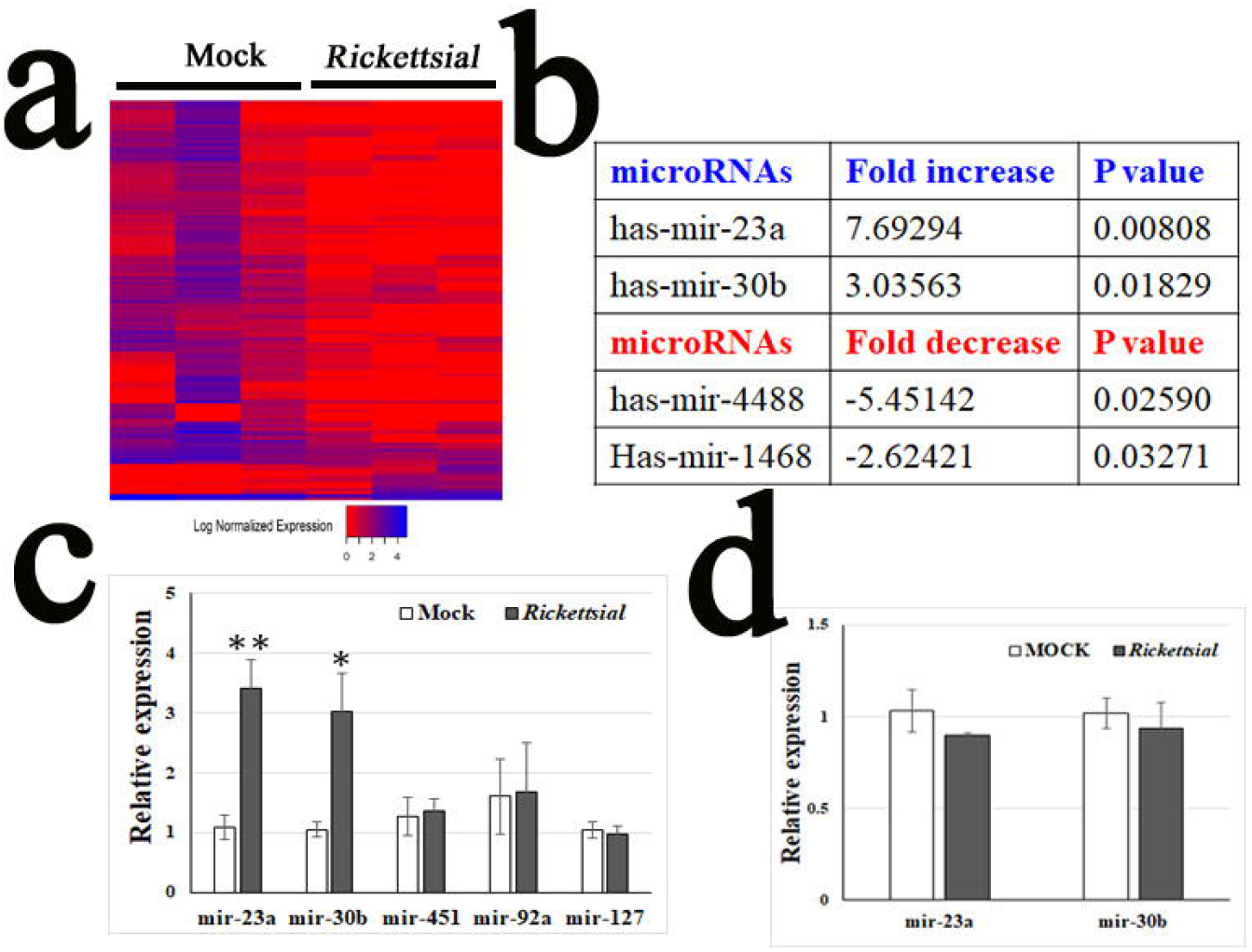
Rickettsial infection alters microRNA expression in ECExos. (**a**) Heatmap clustering of microRNAs in normal ECExos vs. *R*-ECExos (n=3). (**b**) microRNA expression in *R*-ECExos vs. normal ECExos (n=3). (**c**) Stem-loop RT-qPCR analysis of microRNAs obtained from normal ECExos (mock) and *R*-ECExos (rickettsial). ** P < 0.01, * P < 0.05. (**d**) Stem-loop RT-qPCR analysis of microRNAs obtained from normal (mock) and *R*-infected donor HUVECs.

We next validated the enhanced expression of mir-23a and mir-30b in Exos using stem-loop RT-qPCR, which is a common method for detecting sncRNAs in EVs (52, 61, 96, 97). In **Fig. 5c**, exosomal mir-23a was up-regulated after *R* infection with a 3-fold increase in expression compared to the mock group (P < 0.01). Similarly, mir-30b had a near 3-fold increase (P < 0.05). However, the levels of mir-127 (98), mir-451, and mir-92a were stable between mock ECExos and *R*-ECExos (**Fig. 5c**). Furthermore, we did not detect different levels of these miRNAs in cell samples between groups (**Fig. 5d**).

Collectively, our data suggest that miR30b and miR23a are selectively sorted into *R*-ECExos following *R* infection.

## Discussion

ECs are the primary mammalian host target cells of SFR infection (2,5,6). The most prominent pathophysiological effect during SFR infections is increased microvascular permeability, followed by vasogenic cerebral edema and non-cardiogenic pulmonary edema with potentially fatal outcomes (2,5). Cellular and molecular mechanisms underlying endothelial barrier dysfunction in rickettsiosis remains largely unknown (7-9). The novel findings in the present study are that *R* infection increases heterogeneous plsExos release, but endothelial Exo size, morphology, and production are not significantly altered following infection. Mouse *R*-plsExos induced dysfunction of normal mouse recipient BMECs in a dose-dependent manner, and human R-ECExos induced dysfunction of normal human recipient BMECs in an exosomal RNA cargo-dependent manner. Next-generation sequencing and stem loop RT-qPCR analyses suggest that mir-23a and mir-30b are selectively sorted into *R*-ECExos after *R* infection. To our knowledge, this is the first report of studying EVs in the context of obligately intracellular bacterial infections.

Exos are in a similar size range as viruses (33, 94) and contain many types of biomolecules, including proteins and nucleic acids that contribute to diseases pathogenesis, and are being actively investigated in cancers, as biomarkers, and as potential therapeutics (33-38, 68, 70). Exos have been studied in the context of different infections (33-38, 70, 71). During infection, EVs released from the host can be derived from the pathogen or the host. It has been reported that pathogens can utilize different mechanisms to hijack host Exos to maintain their survival and increase their pathogenicity (33). *Mycobacterium tuberculosis* releases lipoarabinomannan into Exos to decrease the interferon response of the recipient macrophage (33). Exosomal gp63 from leishmania has been shown to downregulate the proinflammatory genes in dendritic cells and macrophages (99). Exos released from *Leishmania donovani*-infected macrophages block the formation of microRNA-122 in recipient hepatocytes, resulting in a higher pathogen burden (100). However, most enveloped virions are the same size as Exos, and major exosomal surface markers CD63 and CD81 are enriched in enveloped viruses (54). Such similarities make the separation of virions and Exos in infected samples particularly challenging (54). *Rickettsia* are strictly intracellular bacteria whose size is about 2.0 µm in length (22, 101, 102). Taking advantage of the SEC technology, we succeeded in isolating and purifying bacteria-free plsExos and ECExos from *R*-infected mouse plasma and cell culture media, respectively, having laid the technical foundation for studying the potential role of Exos in the pathogenesis of rickettsiosis.

Differential centrifugation has been employed for Exo isolation for many years, but the technique suffers from aggregation and decreased integrity of Exos (38, 53, 64-66). Recently, single-step SEC was employed successfully for Exo purification with improved integrity, yield, and no aggregation (38, 53, 64-66). In the present study, using SEC technology, we have successfully isolated Exos from plasma and culture media in BSL-2/3 facilities and validated Exo quality using multiple EV-specific assays (**Fig. 1**), demonstrating size-purity and morphologic integrity without aggregation. Exo size and morphology were not significantly changed after *R* infection (**Fig. 1**). The generation of plsExos was significantly upregulated after *R* infection, while no difference was detected in plasma protein concentrations. However, our *in vitro* endothelial *R* infection model demonstrated no significant difference in exosomal generation between normal and *R*-infected ECs. Circulating Exos have been identified as heterogeneous and derived from multiple different types of cells, including ECs, epithelial cells, leukocytes, erythrocytes, and platelets (90). Our data suggest that endothelial Exo size, morphology, and production were not significantly altered after infection.

Exos can induce functional responses by multiple mechanisms, including releasing bioactive components after internalization (67, 68). In infectious disease biology, EVs from infected donor cells are associated with virulence of the pathogens in recipient cells (33-38, 69-71). In the present study, both *R*-plsExos and *R*-ECExos weakened the barrier function of the normal mouse ECs. Concomitantly, human *R*-ECExos induced disruption of the tight junctional protein ZO-1 in recipient human BMECs in an exosomal RNA-dependent manner. However, the underlying mechanism remains unclear.

The discovery of extracellular sncRNAs in the blood, despite the abundant presence of RNases, led to the proposal of a scenario in which sncRNAs are encapsulated in EVs (55, 72-74) or in the form of circulating ribonucleoproteins (75, 76). EV-enclosed messenger RNAs are mostly fragmented, and extracellular RNAs are enriched in sncRNAs (77). Despite a previous report that the average copy number of miRNAs in each EV is low (103), accumulating evidence suggests a critical function of EV-containing miRNAs. EV RNA cargo mostly consists of sncRNAs (77). A growing number of reports have established that many, if not all, the effects of EVs are mediated by microRNAs (52, 55-60, 63), which remain functional to regulate cellular behaviors of the recipient cell (79). Exosomal microRNAs are of particular interest due to their participation in posttranslational regulation of gene expression. A single microRNA can regulate many target genes to affect biological function (104). Recent studies provide emerging evidence that microRNAs are selectively sorted into EVs, independent of their cellular levels (52, 55-62). We observed no significant differences in total sncRNAs (<150 nucleotides) per Exo between normal ECExo and *R*-ECExo. Seventy-two hours after *R* infection, expression levels of mir-23a and mir-30b were remarkably upregulated in *R*-ECExos, but no change was observed in the mock-infected cells. These data suggest that miR30b and miR23a are selectively sorted into *R*-ECExos during *R* infection. The underlying mechanism is yet to be elucidated.

Analysis of the interactions among enriched exosomal microRNAs and potential mRNA targets will provide putative mRNA candidates for future studies. Given that the molecular and functional effects of mir-23a (105-108) and mir-30b (108, 109) have been documented to target endothelial barrier functions, further research into the selective sorting mechanism(s) and functional roles of exosomal mir-23a and mir-30b may provide new insights into the pathogeneses of SFR. Furthermore, additional research may validate specific exosomal microRNAs as impactful druggable targets for the prevention and treatment of fatal human diseases caused by *Rickettisia* and other pathogens.

## Materials and Methods

### Mouse model of *R. australis* infection

All animal experiments were performed according to protocols approved by the Institutional Animal Care and Use Committee of the University of Texas Medical Branch (UTMB). Wild-type (WT) mice (C57BL/6J) were obtained from Jackson Laboratory (Bar Harbor, ME). All mice used in this study were 8 to 12 week-old males. C57BL/6J mice are highly susceptible to *R. australis*. Therefore, this organism was chosen as the SFG rickettsial agent of choice (10). The male C57BL/6 mouse–*R. australis* model is an established animal model of human SFG rickettsiosis because the pathology involves disseminated endothelial infection and pathological lesions, including vasculitis in multiple organs, similar to what is observed in human SFG rickettsiosis (10, 87). After an ordinarily lethal dose of 2 LD_50_ *R. australis* (the LD_50_ is 1 x 10^6^ PFU) was injected through the tail vein (87), blood samples were collect on day 4 p.i. for plasma samples.

### Nanoparticle tracking analysis (NTA)

NTA was performed to determine the size and concentration of EVs at Nanomedicines Characterization Core Facility (The University of North Carolina at Chapel Hill, Chapel Hill, NC). Briefly, isolated Exo samples were diluted to a concentration of 5×10^9^ to 1×10^11^ particles/ml in filtered PBS. The samples were then run on a NanoSight NS500 (NanoSight, Malvern Instruments, Westborough, MA) to capture particles moving by way of Brownian motion (camera type, sCMOS; camera level, 16; detection threshold, 5). The hydrodynamic diameters were calculated using the Stokes-Einstein equation. The 100-nm standard particles and the diluent PBS alone were used for reference.

### microRNA quantification in HUVEC Exos

As previously reported (52), RNAs were extracted from purified HUVEC Exos. Small RNAs (6–150 nucleotides) and microRNA fractions (10–40 nucleotides) were quantified using high-resolution small RNA analysis (Agilent 2100 Bioanalyzer system, Santa Clara, CA) at the Biopolymer Facility (Harvard Medical School, Cambridge, MA). To determine the concentration of small RNAs and microRNAs per Exo, the quantified sncRNA/microRNA value was normalized to the Exo count, which was evaluated using NTA.

### Bioinformatic analysis of sequencing data

Sequencing was done using an Illumina NextSeq as single-end 75 base pair reads generating between 4.8 and 40.9 million reads per sample. Quality control of the samples was performed using QIAGEN CLC Genomics Workbench 20.0. Raw sequencing reads were trimmed to remove QIAGEN 3′-AACTGTAGGCACCATCAAT and 5′-GTTCAGAGTTCTACAGTCCGACGATC adapters, as well as filtered based on initial quality assessment. Reads dominated by low-quality base calls, and longer than 55 nucleotides, were excluded from the downstream analyses. Filtered data undergo further RNA-seq analysis using the CLC Genomics Workbench 20.0 RNA-Seq Analysis 2.2 module with RNAcentral noncoding Human RNA (110, 111)(downloaded April 16, 2020), miRbase 22.1, and the ENSMBL GRCh38 noncoding RNA gene collection (112)(downloaded November 20, 2019). Differential expression analysis was performed using the “Differential Expression in Two Groups 1.1” module. The differential expression module uses multi-factorial statistics based on a negative binomial generalized linear model (GLM) to correct for differences in library size between the samples and the effects of confounding factors. The Wald test was used to compare the expression of noncoding RNA between the groups.

### Atomic force microscopy (AFM)

The purified and concentrated EV sample was diluted at 1:10, 1:100, and 1:1000 with molecular grade water. Glass coverslip were cleaned three times with ethanol and acetone, then three times with molecular grade water. The coverslip was correctly labeled and placed in the hood to dry under laminar flow for an hr and subjected to coating with the diluted EV samples on the designated area for 30 minutes. EV samples were washed away gently with molecular grade water, and the coverslip was dried for one hr.

The coverslip coated with EV samples was examined using an AFM (CoreAFM, Nanosurf AG, Liestal, Switzerland) using contact mode in the air. A PPP-FMR-50 probe (0.5-9.5N/m, 225µm in length and 28µm in width, Nanosensors) was used. The parameters of the cantilever were calibrated using the default script from the CoreAFM program using the Sader *et al*. method. (113) The cantilever was approached to the sample under the setpoint of 20 (113)nN, and topography scanning was done using the following parameters: 256 points per line, 1.5 seconds per line in a 5-µm x 5-µm image.

### Rickettsiae, cell culture, and *R. parkeri* infection

*R. australis* (Cutlack strain) (87) and *R. australis* (Atlantic rainforest strain)(114) were prepared as described. Uninfected Vero cells were processed as mock control material using the same procedure. All biosafety level (BSL)-3 or ABSL-3 experiments were performed in CDC-certified facilities in the Galveston National Laboratory at UTMB, Galveston, TX, using established procedures.

A standard protocol to isolate brain microvascular endothelial cells (BMECs) from wild-type mice (115) was used. Human umbilical vein endothelial cells (HUVECs) (Cell Applications, Inc.) or BMECs were cultivated in 5% CO_2_ at 37°C on type I rat-tail collagen-coated round glass coverslips (12 mm diameter, Ted Pella, Redding, CA) until 90% confluence. HUVECs were infected with *R. parkeri* at an MOI of 10. Uninfected ECs were used as mock controls and were subjected to the same procedure. All experiments were performed in triplicate. Normal mouse or rabbit IgGs were used as negative controls.

### ECExos and plsExos isolation, concentration, and permeabilization

#### ECExos isolation and concentration

Donor HUVECs in T75 flasks were infected using 10 MOI of *R. parkeri* or mock-infected for 72 hours, and 11 mL of media was collected. The media were passed through 0.2-µm syringe filters twice. Following the instruction of the manufacturer, 10 mL of filtered media was subjected to the qEV10 column (Izon, New Zealand) for SEC isolation. The number 7 to 10 fractions were collected as the Exo-enriched fractions, which were concentrated using 100,000 MWCO PES Vivaspin centrifugal filters (Thermo Fisher Scientific). Exo samples (in 200 µl PBS) were stored at −80□ prior to use in downstream assays.

#### plsExos isolation and concentration

Blood samples were collected in anticoagulation tubes on day 4 p.i. for plasma isolation. The plasma sample (200 µl) was passed through 0.2µm syringe filters twice. Following the instruction of the manufacturer, filtered plasma was subjected to the qEV10 (Izon, New Zealand)for SEC isolation. The number 7 to 10 fractions were collected as the Exo-enriched fractions, which were concentrated using 100,000 MWCO PES Vivaspin centrifugal filters (Thermo Fisher Scientific). Exo samples (in 200 µl PBS) were stored at −80□ prior to use in downstream assays.

For saponin-assisted active exosomal permeabilization pretreatment (81-83) of Exos using RNase, Exo samples (1 x 10^9^ particles/mL) and RNase (20 µg/mL) (Thermo Fisher Scientific) were incubated with 0.1 mg/ml saponin (Thermo Fisher Scientific) at room temperature for 15 min. After rinsing using phosphate-buffered saline (PBS), Exo samples were concentrated using 100,000 MWCO PES Vivaspin centrifugal filters.

### The distribution of Exos *in vivo* and *in vitro*

Using a published approach, recipient cell uptake of Exos was assessed *in vivo* and *in vitro* (90). Briefly, following incubation using materials from the PKH26 Red Fluorescent Cell Linker Kit (Millipore Sigma, St. Louis, MO), the Exos were washed three times with PBS before ultracentrifugation at 100,000□x□*g* for 20□min at 4□°C using Beckman L7-80 and rotor SW41 (Beckman Coulter, Indianapolis, IN) to remove unbound stain. PBS without Exo was processed with same steps as the mock PKH26-labeled tracer. A single injection of PKH26-labelled exosomes (about 1□x□10^11^ particles in 100□µL of PBS) via the tail vein of a normal mouse was done to observe the distribution of Exos in the lungs, liver, and brain 6□hr after injection. Immunofluorescence staining was done using frozen sections with rabbit antibodies to CD31. For *in vitro* assessment, PKH26-labeled ECExos (2000 particles/cell) were added in the culture media of normal human BMECs. After 2 hrs, cells were fixed. All solutions of PKH26-labeled ECExos were filtered with a 0.2 µm filter. Fluorescent images were analyzed using Olympus BX51 epifluorescence and Nikon A1R MP ECLIPSE T*i* confocal microscope with *NIS*-Elements imaging software version 4.50.00 (Nikon, Tokyo, Japan).

### Stem-loop real time PCR

Total RNA was extracted from EVs by using Trizol (Invitrogen). An exogenous synthetic microRNA, namely cel-mir-39, was diluted in TRIzol before extraction to act as a normalizer. The concentration of total RNA was measured by NanoDrop (ND-2000). TaqMan MicroRNA Reverse Transcription kit (Applied Biosystems) was used for reverse transcription reactions. The 15 ul RT reactions contained 5 ng total RNA template, 3 ul RT Primer (5×), 0.15 ul dNTPs (100 mM), 1 ul MultiScribe reverse transcriptase (50 U/µL), 1.5 ul Reverse Transcription Buffer (10×), 0.19 ul RNase inhibitor (20 U/µL), and 4.16 ul nuclease-free water. Reverse transcription conditions were 16□ for 30 min, 42□ for 30 min, and 85□ for 5 min. For PCR amplification, the 10 ul PCR reactions included 0.7 ul cDNA template acquired above, 0.5 ul TaqMan Small RNA Assay Mix (20X), 5 ul PCR Master Mix, and 3.8 ul nuclease-free water. qPCR reaction conditions were 50□ for 2 min, 95□ for 30 sec, followed by 40 cycles of 95□ for 5 sec, and 65□ for 30 sec. The relative expression of each miRNA was expressed as 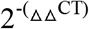 by the CFX Connect Real-Time System (Bio-Rad, Hercules, CA).

### Statistics

Statistical significance was determined using Student’s *t*-test or one-way analysis of variance. Results were regarded as significant if two-tailed P values were < 0.05. All data are expressed as mean ± standard error of the mean.

## Acknowledgements

We gratefully acknowledge Mr. Pragnesh Patel for his contributions establishing the capacity of the exosomal size distribution analysis. We gratefully acknowledge Dr. Kimberly Schuenke for her critical review and editing of the manuscript. We thank Dr. Hugo Samano for input during the planning phases of the experiments. This work was supported by NIH grant R01AI121012 (BG), R21AI137785 (BG), R21AI154211(BG), R03AI142406 (TS and BG), and R21AI144328 (TS and BG). The sponsors had no role in the study design, data collection and analysis, decision to publish, or preparation of the manuscript.

## Authorship Contributions

BG and TS designed the study, performed experiments, analyzed data, and wrote the manuscript. YL, CZ, ZS, QC, KK performed experiments, analyzed data, and the prepared manuscript. YQ, JB, AG, JX and AD analyzed data. YJ and GG designed the study and analyzed data.

