## supplementary materials for "Endothelial exosome plays functional role during rickettsial infection"

**Materials and Methods**

**Antibodies and other reagents**

Anti-AnnexinA2 mouse monoclonal (mAb) antibodies (clone 666316) were purchased from R&D Systems (Minneapolis, MN) via Thermo Fisher Scientific (Rockford, IL). Anti-VE-cadherin rabbit antibody, anti-CD31 rabbit antibody, anti-CD45 and anti-CD63 rabbit antibody were purchased from Abclonal (Woburn, MA). Anti-flotillin-1 mouse antibody and anti-calnexin mouse antibody were purchased from BD Transduction Laboratories (San Jose, CA). Anti TSG101 rabbit antibody was purchased from Novus Biologicals (Centennial, CO). Anti-albumin rabbit antibody was purchased from cell signal technology (Beverly, MA). Anti-von Willebrand rabbit polyclonal antibody, anti-ZO-1 rabbit antibody, AlexaFluor 488-conjugated goat anti-mouse IgG, AlexaFluor 594-conjugated goat anti-rabbit IgG, and DAPI were purchased from Invitrogen (Carlsbad, CA). Normal mouse and rabbit IgGs were purchased from Agilent (Santa Clara, CA). Endothelial Cell Growth Medium and fetal bovine serum were obtained from Cell Applications, Inc. (San Diego, CA). MicroBCA protein assay kit was purchased from Thermo Fisher Scientific (Rockford, IL). Unless otherwise indicated, all reagents were purchased from Thermo Fisher Scientific.

**Transmission electron microscopy** (**TEM)**

Negative staining was used to visualized EVs under TEM. A TEM grid was coated with 0.01% bovine serum albumin (BSA) for 30 seconds. Then the BSA was drawn off from the TEM grid and 10 ul of EV sample was immediately added and left on for one minute. Then the grid was washed with 1mM ethylenediaminetetraacetic acid (EDTA) and stained with 10 ul 0.5% uranyl acetate, which was drawn off from the edge of the grid after one minute. The TEM grid was dried under a heat lamp for three minutes before observation under TEM. All the solutions were filtered using 0.2µm syringe filters.

**Western immunoblotting**

For Western immunoblotting, equal amounts of soluble protein were subjected to 10% SDS–polyacrylamide gel electrophoresis (SDS-PAGE). Proteins were transferred onto a polyvinylidene difluoride membrane and then incubated with primary antibody (1:1,000 for anti- Flotillin-1, Albumin, AnnexinA2, CD63, TSG101, and calnexin-1 antibodies) at 4℃ overnight, followed by incubation with a secondary antibody at 1:10,000 for 2 hrs. A goat anti-mouse or rabbit IgG and IgM (H+L)-HRP (Thermo Fisher Scientific) were used as the secondary antibodies. Blots were visualized using Pierce™ ECL Western Blotting Substrate kit (Thermo Fisher Scientific).

**Trans-endothelial electrical resistance (TEER)**

TEER was measured using an EVOM resistance meter (Millicell ERS-2) (Thermo Fisher Scientific, Rockford, IL), as reported (1). TEER of the monolayer of BMECs seeded on inserts in 24-well plates (0.4 µm polyester membrane, CoStar, Thermo Fisher Scientific) were measured after 48 h of treatment. The values were shown as Ω×cm^2^ and normalized by subtracting the background (TEER from an insert without cells).

**Immunofluorescence (IF)**

For IF staining of vWF in mouse tissues collected after extensive *in vivo* perfusion, Ultra V Block and normal rabbit serum were employed for the reduction of nonspecific background before and after vWF antigen, respectively. Frozen sections of 5 µm thickness were incubated with anti-vWF rabbit polyclonal antibody (1:500) for 2 hrs at room temperature, followed by AlexaFluor 488-conjugated goat anti-rabbit IgG (1:1000) for 30 minutes at room temperature. For IF of VE-cadherin or ZO-1 in BMECs, cells were incubated with anti-VE-cadherin or ZO-1 rabbit polyclonal antibody (1:500) for 2 hours, followed by AlexaFluor 594-conjugated goat anti-rabbit IgG (1:1000) for 30 minutes. Nuclei were counter-stained with DAPI. A rabbit polyclonal IgG (Thermo Fisher) served as a negative control (2) (**Supple Fig. 2**). Fluorescent images were analyzed using an Olympus BX51 epifluorescence or Nikon A1R MP ECLIPSE T*i* confocal microscope with *NIS*-Elements imaging software version 4.50.00.

**Supple Fig1**: The quantities of rickettsiae in plasma-derived Exos (plsExos) (n=5/group) and HUVEC culture media-derived Exos (ECExos) (n=3/group) were determined by quantitative real time PCR. Data are presented as means ± standard errors.

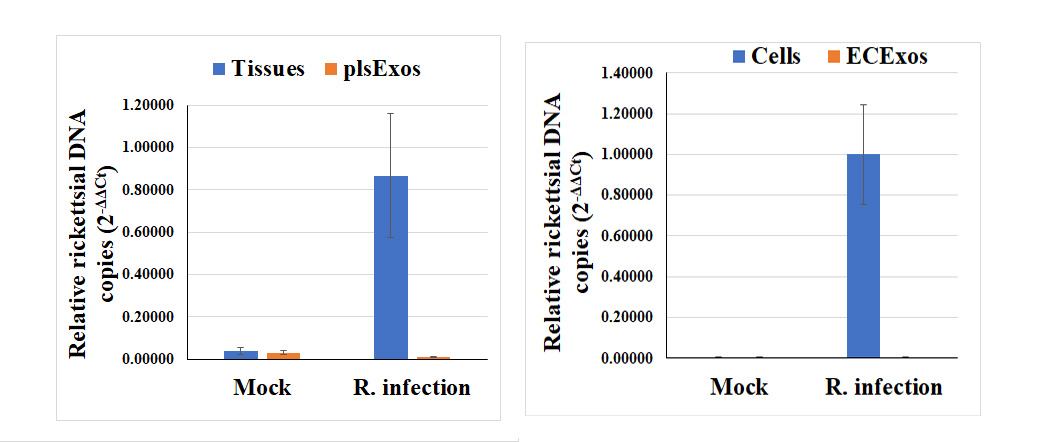

**Supple Fig 2**: Normal rabbit serum was used as an antibody during IF in tissue (a) and cells (b). Scale bar indicates 20 µm.

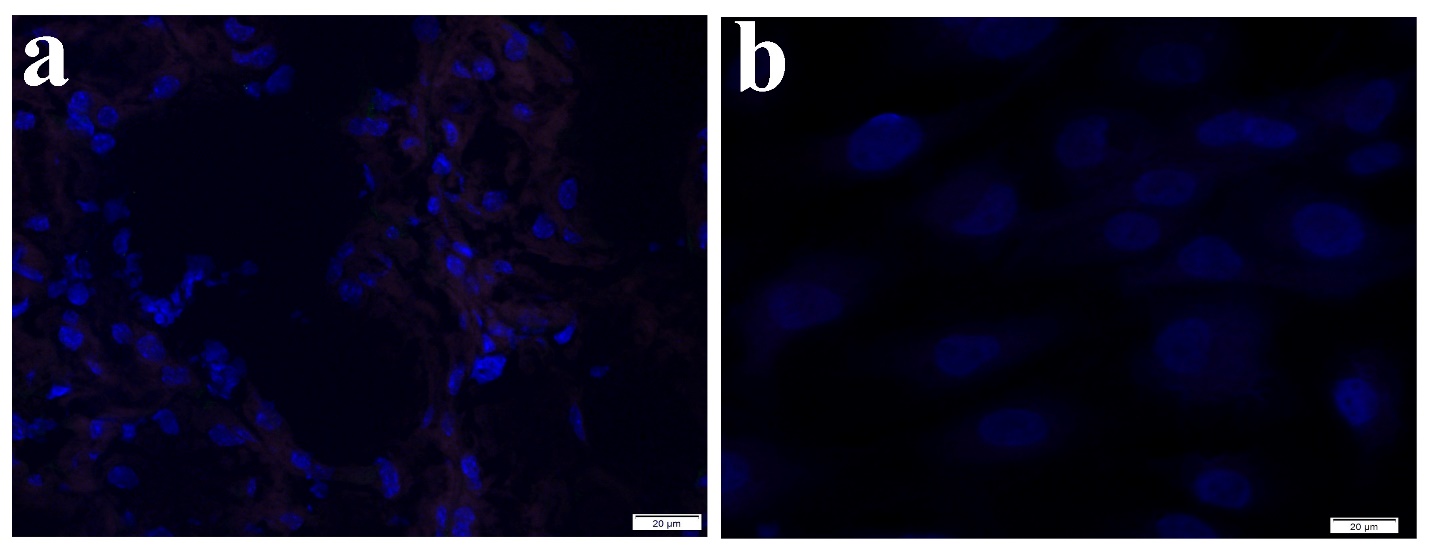

1.	Wilhelm I, Fazakas C, Krizbai IA. 2011. In vitro models of the blood-brain barrier. Acta Neurobiol Exp (Wars) 71:113-28.

2.	Liu Y, Xiao J, Zhang B, Shelite TR, Su Z, Chang Q, Judy B, Li X, Drelich A, Bei J, Zhou Y, Zheng J, Jin Y, Rossi SL, Tang SJ, Wakamiya M, Saito T, Ksiazek T, Kaphalia B, Gong B. 2020. Increased talin-vinculin spatial proximities in livers in response to spotted fever group rickettsial and Ebola virus infections. Lab Invest doi:10.1038/s41374-020-0420-9.
